# Mitophagy mediates beige to white transition of human primary subcutaneous adipocytes *ex vivo*

**DOI:** 10.1101/2022.02.01.478645

**Authors:** Attila Vámos, Abhirup Shaw, Klára Varga, István Csomós, Gábor Mocsár, Zoltán Balajthy, Cecília Lányi, Zsolt Bacso, Mária Szatmári-Tóth, Endre Kristóf

## Abstract

Brown and beige adipocytes have multilocular lipid droplets, express uncoupling protein (UCP) 1, and promote energy expenditure. In rodents, when the stimulus of browning subsides, parkin-dependent mitophagy is activated and dormant beige adipocytes persist. In humans, however, the molecular events during beige to white transition were not studied in detail. In this study, human primary subcutaneous abdominal preadipocytes were differentiated to beige for 14 days, then either the beige culture conditions were applied for additional 14 days or it was replaced by a white medium. Control white adipocytes were differentiated by their specific cocktail for 28 days. Peroxisome proliferator-activated receptor γ-driven beige differentiation resulted in increased mitochondrial biogenesis, UCP1 expression, fragmentation, and respiration as compared to white. Morphology, UCP1-content, mitochondrial fragmentation, and basal respiration of the adipocytes that underwent transition, along with the induction of mitophagy, were similar to control white adipocytes. However, white converted beige adipocytes had a stronger responsiveness to dibutyril-cAMP, which mimics adrenergic stimulus, than the control white ones. Gene expression patterns showed that removal of mitochondria in transitioning adipocytes may involve both parkin-dependent and independent pathways. Preventing entry of beige adipocytes into white transition can be a feasible way to maintain elevated thermogenesis and energy expenditure.

## 1. Introduction

The prevalence of obesity has dramatically increased worldwide in the last decades [1], which can be linked with many factors, including metabolic, genetic, environmental, and behavioral impacts [2,3]. Obesity significantly enhances the risk of many diseases, including metabolic syndrome, type 2 diabetes, non-alcoholic fatty liver disease, cardiovascular diseases, or certain types of tumors, which are leading causes of death today [4-7]. Recently, obesity was recognized as an independent risk factor for mortality from severe acute respiratory syndrome coronavirus 2 infections [8,9]. The available treatments are interventions of diet, exercise, or lifestyle and bariatric surgery [10,11]; however, effective anti-obesity therapeutic strategies are strongly limited.

Brown adipose tissue (BAT), which contains thermogenic brown and beige adipocytes, is capable for energy dissipation through the production of heat, mainly mediated by Uncoupling protein (UCP) 1-dependent proton leak that uncouples oxidative phosphorylation from ATP generation in the mitochondria [12-14].BAT plays a key role in maintaining the constant core body temperature through non-shivering thermogenesis and might open up promising therapeutic opportunities to combat obesity [15,16]. BAT can be found in six anatomical regions (cervical, supraclavicular, axillary, mediastinal, paraspinal, and abdominal) in adult humans, which amounted to 4.3% of total fat and 1.5% of total body mass [17]. Based on mathematical predictions, BAT can oxidize around 4 kg fat per a year in adult humans and its thermogenic activity can contribute up to 5% of the basal metabolic rate [18].

White adipocytes function as long-term energy storage, contain few mitochondria, a single large lipid droplet, and express low level of UCP1 [19,20]. Brown and beige adipocytes originate from distinct precursor cells, characterized by multilocular small lipid droplets, high mitochondrial content, and detectable level of UCP1 expression [21,22]. Beige and white adipocytes are derived from the same mesenchymal precursors and the beige cells can be found in a masked condition in subcutaneous white adipose tissue (WAT) depots [21]. Beige adipocytes contain low amount of UCP1 in basal conditions, and have to be activated, e.g. by cold, for thermogenesis, which is mainly mediated through β-adrenergic stimulation [23]. However, the regulation of beige adipocyte maintenance and inducibility (which process is often called “browning”) in humans has remained mainly elusive.

Autophagy is a well-described intracellular catabolic process, in which cargoes, such as protein aggregates or damaged organelles, are delivered by double-membrane-bound structures, termed autophagosomes, to the lysosomes for degradation and their components are recycled [24,25]. Many highly conserved autophagy-related (ATG) proteins are responsible for the biogenesis of autophagosome [26]. Upon induction of autophagy, the cytosolic form of Microtubule-associated protein 1A/1B-light chain 3 (LC3)-I is conjugated to phosphatidylethanolamine to generate the lipidated LC3-II, which is then recruited to autophagosomal membranes. LC3 is a well-accepted autophagosome marker, since LC3-II-content is an indicator of the amount of autophagosome formation [27]. Detecting the conversion of LC3-I to LC3-II by immunoblotting is a commonly used method to follow autophagy activity [28,29].

Damaged or unwanted mitochondria can be removed by selective autophagy, termed mitophagy, which is considered as a crucial mechanism of mitochondrial quality control [30,31]. In the adapter-mediated, ubiquitin-dependent mitophagy pathway, mitochondrial depolarization initiates the accumulation of phosphatase and tensin homolog–induced putative kinase (PINK) 1 in the outer mitochondrial membrane, resulting in the recruitment of parkin from the cytosol, which ubiquitinates the outer mitochondrial proteins. The selective autophagy adapter proteins, such as, Calcium Binding And Coiled-Coil Domain 2/Nuclear Domain 10 Protein 52 (CALCOCO2/NDP52), Optineurin (OPTN), Neighbor Of BRCA1 Gene (NBR) 1, and p62 (encoded by SQSTM1 gene) link the parkin ubiquitinated mitochondrial proteins and LC3, leading to the sequestration of mitochondria into autophagosomes [32,33]. Furthermore, the adapter-independent, ubiquitin-independent mitophagy process is characterized by the direct interaction between LC3 and mitochondria-localized proteins, such as BCL2 Interacting Protein (BNIP) 3, BNIP3 Like/NIP3-Like Protein X (BNIP3L/NIX), or FUN14 Domain Containing (FUNDC) 1 [34,35]. We have considerable knowledge about selective mitophagy, however, many questions remain unanswered, such as the extent to which mitochondrial clearance is regulated in a cell type or tissue specific manner.

Recent publications proved the significance of autophagy in the regulation of beige adipocyte thermogenesis [36]. In rodents, Kajimura and his collaborators showed the role of parkin-dependent mitophagy in beige to white adipocyte transition as a result of the removal of β-adrenergic stimulus, which resulted in inactive, but reactivation capable beige adipocytes with white morphology [37,38]. However, this process has not been identified and characterized in humans so far. Recently, our research group described how cyclic adenosine monophosphate(cAMP)-driven thermogenic activation regulates mitophagy in human masked and mature beige adipocytes *ex vivo*. Our data indicated continuous mitochondrial clearance in these adipocytes, which was rapidly repressed in response to short term adrenergic stimulus, pointing to a fast regulatory mechanism to provide high mitochondrial content for thermogenesis [39].

In this study, we have followed up the cell autonomous transition of human primary subcutaneous abdominal beige adipocytes to white and investigated whether mitophagy is activated during this process. The transition, which involved both parkin-dependent and independent pathways, resulted in characteristic gene expression, morphological, and functional features of the white adipocytes; however, the converted cells could be strongly activated by a cell permeable cAMP analogue.

## 2. Results

### 2.1. Thermogenic Competency of Human Abdominal Subcutaneous Derived Adipocytes Is Induced Following Continuous Peroxisome Proliferator-Activated Receptor (PPAR) γ Stimulation and Subsides as a Result of Beige to White Transition

To study the adipogenic potential of primary human adipose-derived stromal cells (hASCs) and the thermogenic competency of differentiated adipocytes, our research group optimized previously published white [40] and brown/beige [41] adipogenic differentiation protocols. These regimens contain diverse compositions of hormones, in which the PPARγ agonist, rosiglitazone is the key driver of browning [42-44]. As expected, abdominal subcutaneous hASCs expressed the major functional marker gene and protein of thermogenesis, UCP1 at the limit of detection. A moderate UCP1 expression was found in adipocytes that were differentiated up to 28 days to white *ex vivo* (Figure 1A, B). Consistent with previous results, the continuous PPARγ stimulation resulted in a marked increase in gene and protein expression of UCP1 in adipocytes differentiated to beige compared with the white ones. UCP1 was further upregulated when beige differentiation was carried out for three or four weeks, respectively. When the beige cocktail was replaced by the white and rosiglitazone was omitted at the fourteenth day of differentiation, UCP1 gene and protein expression tended to elevate for the following week similarly to those adipocytes that were continuously exposed to the beige regimen. After two weeks of rosiglitazone withdrawal, UCP1 gene and protein expression was significantly decreased as compared to the beige adipocytes (Figure 1A,B) and showed a gene expression level that was comparable to white adipocytes (Figure 1A). The decline of UCP1 expression at protein level was slower (Figure 1B) than at mRNA level (Figure 1A).

**Figure 1.**
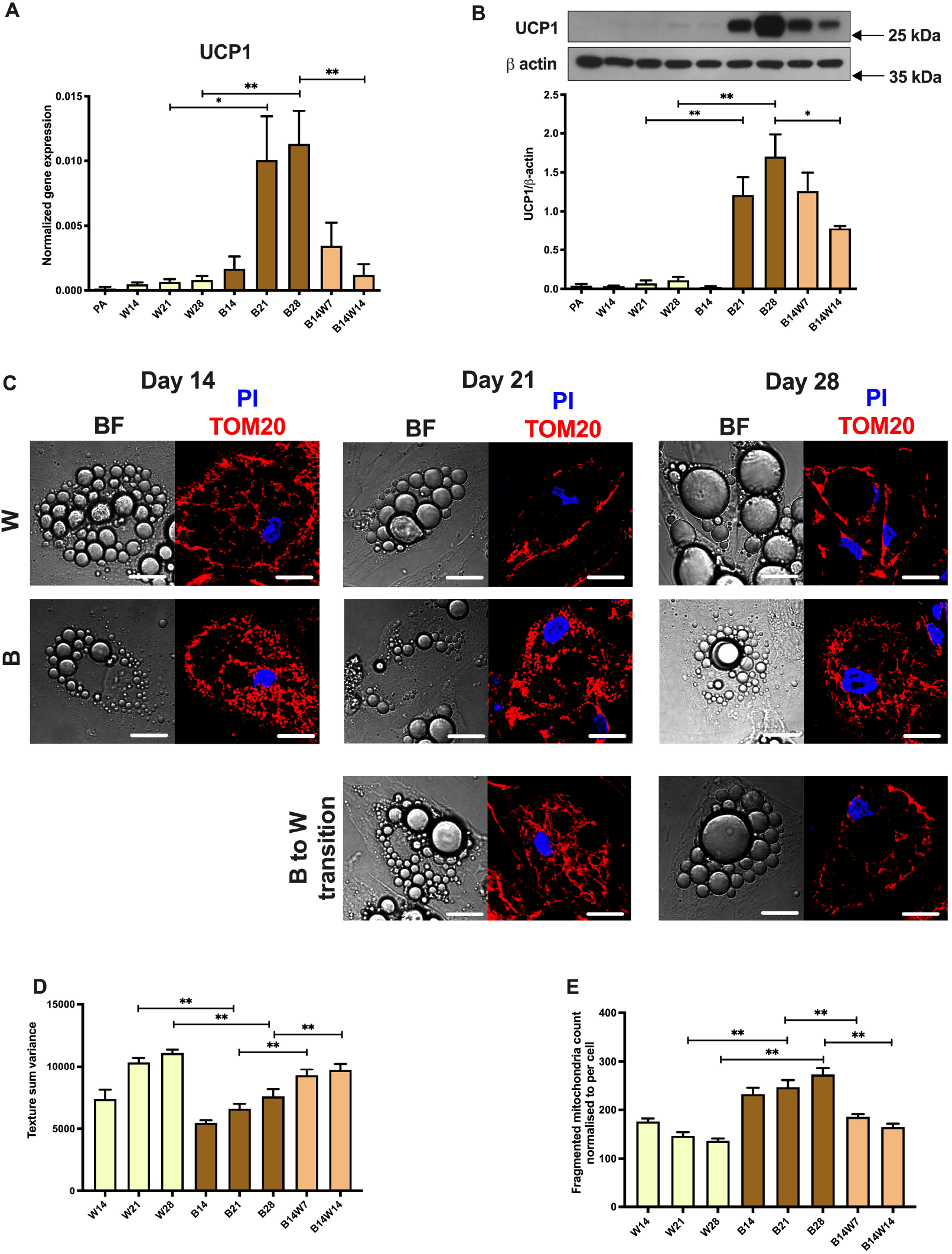
Mitochondrial uncoupling protein (UCP) 1 expression and fragmentation were elevated following long term rosiglitazone treatment and subsided upon beige (B) to white (W) transition. Human primary abdominal subcutaneous preadipocytes were differentiated to B for 14 days (B14), then either the B culture conditions were applied for additional 14 days (B21 and B28) or it was replaced by a W differentiation medium (B14W7 and B14W14). As a negative control, W adipocytes were differentiated by their specific cocktail (W14, W21, and W28). Quantification of UCP1 (A) gene and (B) protein expression (n=6), (C) Representative confocal microscopy images of TOM20 immunostaining, nuclei were labelled with propidium iodide (PI), and BF represents brightfield image, scalebars represent 10 μm, (D) Texture sum variance quantified from BF images (n=50 cells from 3 donors), (E) Fragmented mitochondrial content quantified based on TOM20 immunostaining normalized to per cell (n=50 cells from 3 donors). Gene expression was normalized to GAPDH and protein expression to β-actin. Data are presented as Mean ± SD. *p<0.05, **p<0.01. Statistics: one-way ANOVA with Tukey’s post-test.

As expected, white adipocytes gained a few but large lipid droplets in a time-dependent manner. Contrarily, beige differentiation resulted in numerous and smaller droplets. However, the droplets merged and enlarged when the beige protocol was replaced by the white (Figure 1C). The quantification of this phenomenon by texture sum variance, which value separated well the white and beige adipocyte populations previously in several cellular models [43-46], is shown in Figure 1D. To investigate the mitochondrial morphology, we performed immunostaining for translocase of outer mitochondrial membrane (TOM) 20 [39,47] (Figure 1C; see secondary antibody control in Figure S1, Supplementary Materials). Consistent with the literature, the number of fragmented mitochondria that reflect thermogenic potential [48] was higher during beige compared to white differentiation. However, the amount of fragmented mitochondria was decreased to the same extent that of the white adipocytes within a week when the beige regimen was discontinued and followed by the white (Figure 1C,E). The changes in UCP1-content and morphological features of the adipocytes suggest that they strongly increase their thermogenic competency, even up to four weeks, as a result of PPARγ agonist, however, could undergo beige to white transition in response to the removal of the browning-inducer.

### 2.2. Elevated Mitochondrial Content, Respiration, and Extracellular Acidification of Beige Adipocytes Disappear After Their Transition to White Phenotype

As a next step, we intended to investigate the mitochondrial content and function during the differentiation and transition. As expected, mitochondrial DNA (mtDNA) amount was higher in beige compared to white differentiated adipocytes and was subsequently decreased as a result of beige to white transition (Figure 2A). White and beige differentiation resulted in the same expression level of the mitochondrial biogenesis master regulator, PPARγ coactivator (PGC) 1α [49]. In response to transition, PGC1α tended to be downregulated, however, this effect did not reach the level of statistical significance. Undifferentiated progenitors expressed PGC1α only at the limit of detection (Figure 2B).

**Figure 2.**
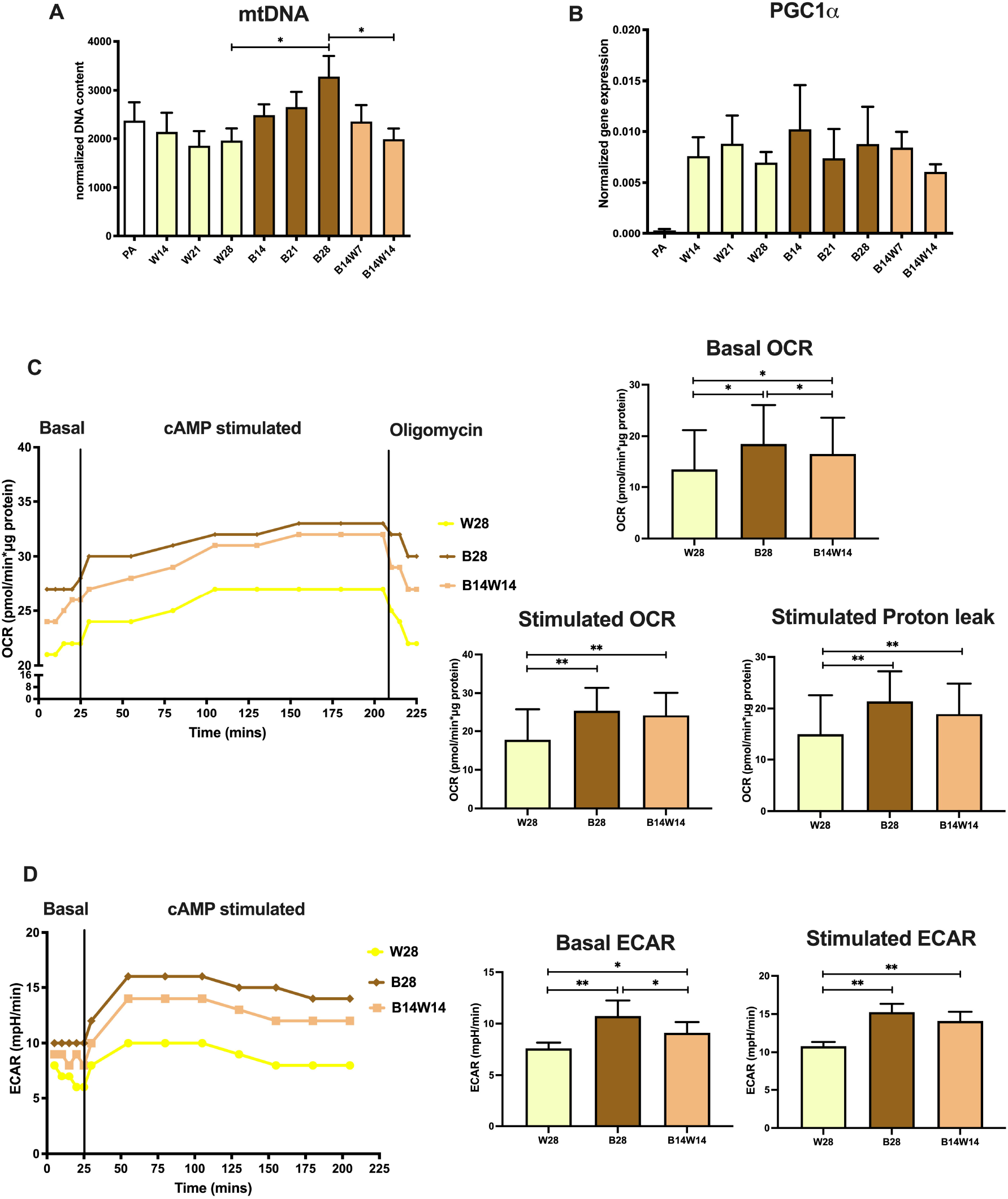
Functional parameters of mitochondrial biogenesis and thermogenesis in white (W), beige (B) adipocytes, and in response to transition. Human primary abdominal subcutaneous preadipocytes were differentiated as in Figure 1. Quantification of (A) total mitochondrial DNA content (n=6) and (B) PGC1α gene expression (n=6), (C) Representative oxygen consumption rate (OCR) curve, followed by quantification of basal, stimulated, and stimulated proton leak OCR (n=4), (D) Representative extracellular acidification rate (ECAR) curve, followed by quantification of basal and stimulated ECAR (n=4). Gene expression was normalized to GAPDH. Data are presented as Mean ± SD. *p<0.05, **p<0.01. Statistics: one-way ANOVA with Tukey’s post-test.

Then we carried out extracellular flux analysis to reveal the functional parameters of differentiated adipocytes [44,50,51]. In parallel with previously reported data [43,51], basal oxygen consumption rate (OCR) of beige adipocytes was higher than of the white ones. Adipocytes undergoing two weeks of transition had significantly higher basal OCR than the white, but lower than the beige adipocytes, which were differentiated for the same period of time. As expected, the cell permeable cAMP analogue, which mimics adrenergic stimulus driven activation of thermogenesis, promptly increased OCR of each type of adipocytes. Proton leak respiration of cAMP stimulated adipocytes, which positively correlates with UCP1 activity, could be assessed after the inhibition of ATP synthase complex by oligomycin [50,52]. Beige adipocytes had elevated stimulated and proton leak OCR compared to the white ones. After the transition, these parameters remained comparable to those observed in beige adipocytes (Figure 2C). Consistent with previous results [44,51], basal extracellular acidification rate (ECAR) was increased in beige compared to white adipocytes, while the transition had a significant suppressing effect on this parameter. Although cAMP stimulated ECAR of each adipocyte type, fully differentiated and converted beige adipocytes showed a greater response to the thermogenic cue than the white ones (Figure 2D). In summary, our data suggest that adipocytes which underwent beige to white transition appear as a cell population with distinct features as compared to the fully differentiated white or beige cells.

### 2.3. Autophagy is Increased at Beige to White Transition

In rodents, it was proved that the withdrawal of cold or β3-adrenergic stimuli activates mitophagy and mediates beige to white transition *in vivo* [37]. Primarily, we investigated the expression of ATG genes, which orchestrate autophagosome formation. In response to transition, ATG5 tended to be upregulated at the second week, however, this difference did not reach the level of statistical significance (Figure 3A). The mRNA expression of ATG7 (Figure 3B) and ATG12 (Figure 3C) was significantly increased at the first week of transition compared to beige adipocytes. The investigated ATG genes were expressed markedly lower in undifferentiated preadipocytes and at a higher, but the same extent in white and beige adipocytes (Figure 3A-C).

**Figure 3.**
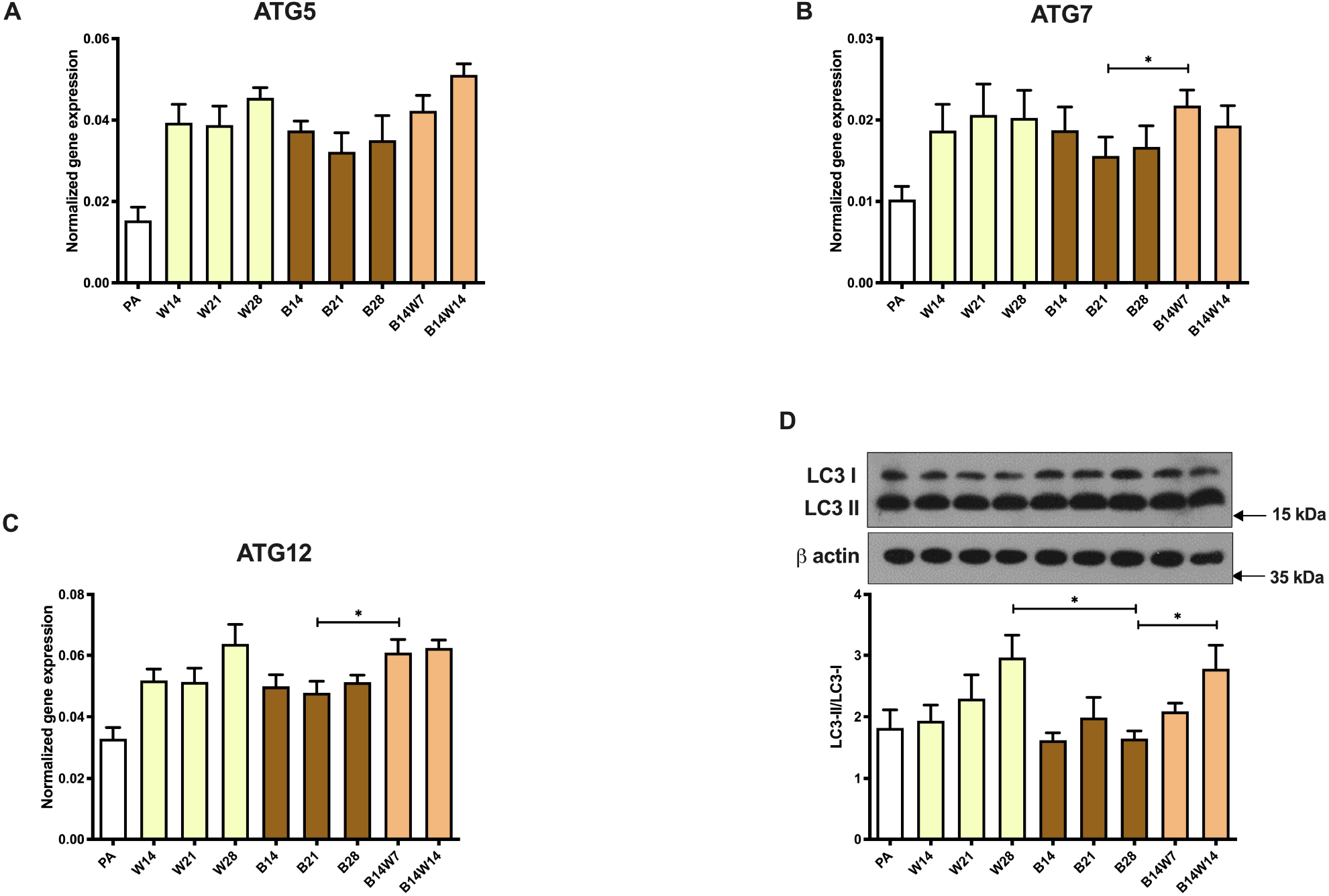
General autophagy markers showed a decreasing trend following long term rosiglitazone treatment and were increased upon beige (B) to white (W) transition. Human primary abdominal subcutaneous preadipocytes were differentiated as in Figures 1 and 2. (A-C) Quantification of ATG5, ATG7, and ATG12 gene expression, (D) representative immunoblot and densitometry analysis of LC3-II/LC3-I protein ratio. Gene expressions were normalized to GAPDH. Data are presented as Mean ± SD. n=6. *p<0.05. Statistics: one-way ANOVA with Tukey’s post-test.

To assess the activity of the ongoing autophagy, we examined the specific autophagy marker, conversion of LC3-I to LC3-II, in adipocytes that were fully differentiated to white or beige or underwent transition. Quantification of this process by immunoblotting is a widely accepted method to monitor autophagy rate [28]. We found a continuous elevation of LC3-II/LC3-I ratio in white adipocytes differentiated up to 28 days, which indicates high autophagy activity. In beige adipocytes, the activity constantly remained at a moderate level, which was significantly lower compared with the white cells, after four weeks of differentiation. When the beige protocol was replaced by the white, autophagy level was significantly increased in two weeks, as compared to the fully differentiated beige adipocytes (Figure 3D). This could be confirmed when the subcellular distribution of LC3 was visualized by immunostaining (Figure 4A; see secondary antibody control in Figure S1, Supplementary Materials). As expected, white adipocytes, which were differentiated to three or four weeks, respectively, contained more LC3 punctae per cell than the beige ones. In addition, the beige to white transition significantly increased the number of LC3 punctae, as compared to the beige adipocytes (Figure 4B). Our data demonstrate that general autophagy was induced in a cell autonomous manner during *ex vivo* beige to white transition of human subcutaneous adipocytes.

**Figure 4.**
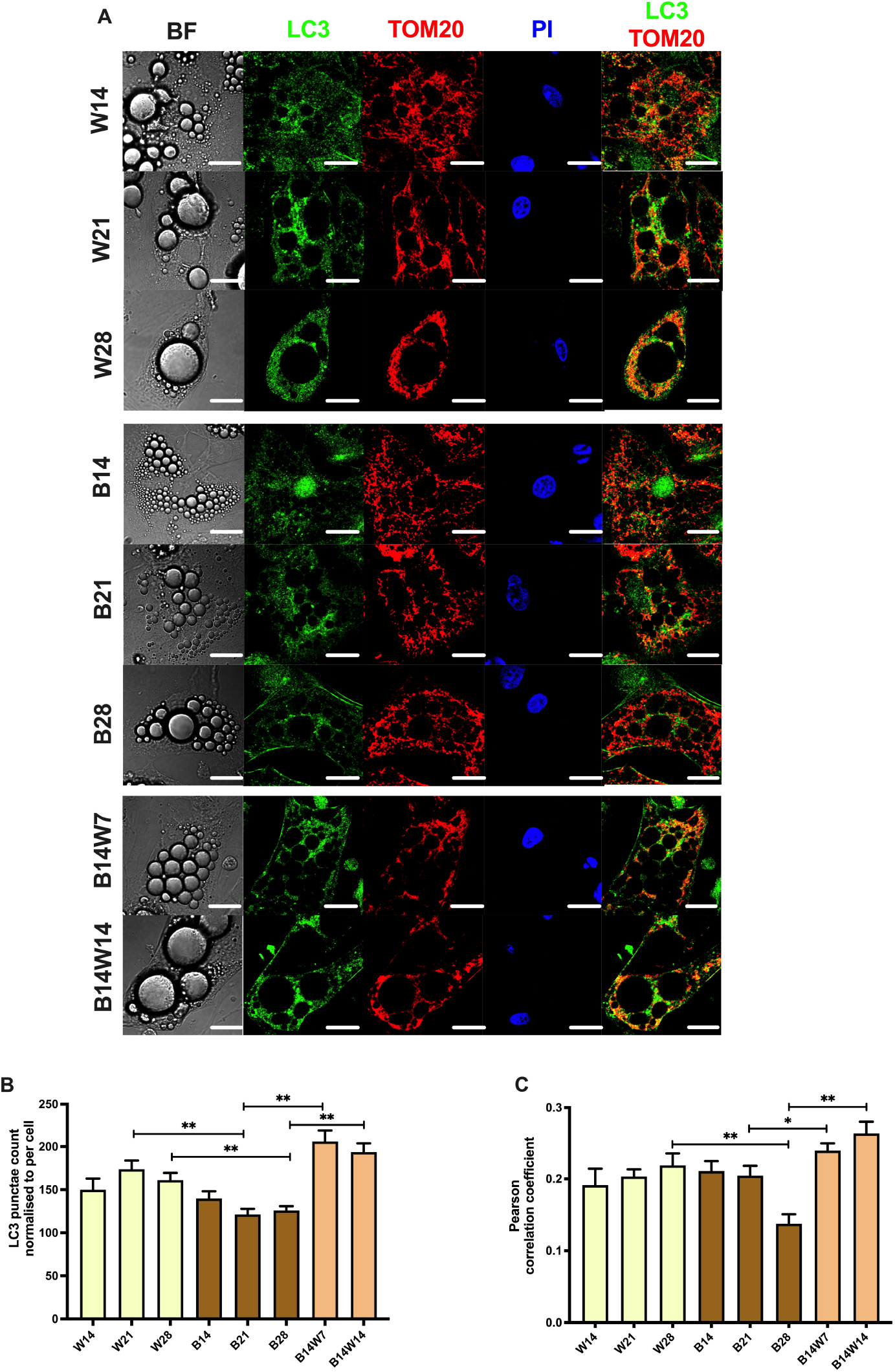
Mitophagy was repressed following long term rosiglitazone treatment and increased upon beige (B) to white (W) transition. Human primary abdominal subcutaneous preadipocytes were differentiated as in Figures 1-3. (A) Representative confocal microscopy images of LC3 and TOM20 immunostaining, nuclei were labelled with propidium iodide (PI), and BF represents brightfield image, scalebars represent 10 μm, (B) Quantification of LC3 punctae normalized to per cell (n=50 cells from 3 donors), (C) Quantification of mitophagy as co-localization of LC3 and TOM20 immunostaining (n=50 cells from 3 donors). Data are presented as Mean ± SD. *p<0.05, **p<0.01. Statistics: one-way ANOVA with Tukey’s post-test.

### 2.4. Beige Differentiation Represses, while Transition to White Increases Mitophagy Involving Selective Autophagy Adapters

We performed co-immunostaining of TOM20 and LC3 (Figure 4A; see secondary antibody control in Figure S1, Supplementary Materials) to follow autophagosome formation and deliverance of mitochondria for degradation [47]. Then we quantified co-localization of the autophagosome and mitochondrial markers by Pearson correlation coefficient (PCC) values. Consistent with the increased autophagy activity (Figure 3), we found elevated PCC values during white adipogenesis of four weeks compared with the beige adipocytes. The co-localization was stronger in adipocytes that underwent transition than in fully differentiated beige cells (Figure 4C).

Parkin, an E3 ubiquitin ligase encoded by the PARK2 gene, is one of the key regulators of mitophagy [53]. Parkin was expressed at a low extent in preadipocytes both at mRNA (Figure 5A) and protein levels (Figure 5B). The applied adipogenic protocols similarly upregulated its expression. Transition did not alter the expression of parkin both at gene (Figure 5A) and protein levels (Figure 5B). Next, we investigated the abundance of selective autophagy adapter proteins which are consumed building a molecular link between the target organelles and LC3-II of the autophagosomes during ongoing mitophagy [54]. NBR1 (Figure 5C) and p62 (Figure 5D) protein content of white adipocytes was comparable to the undifferentiated progenitors and showed a decreasing trend up to 28 days of differentiation. In beige adipocytes, significantly more of the aforementioned adapters could be detected. This further supports low activity of selective autophagy in beige adipocytes. After removal of the browning-inducer from differentiation media, NBR1 (Figure 5C) and p62 (Figure 5D) amount declined significantly in one- or two-weeks periods, respectively. Our data suggest that the selective autophagic degradation of mitochondria is enhanced during beige to white transition.

**Figure 5.**
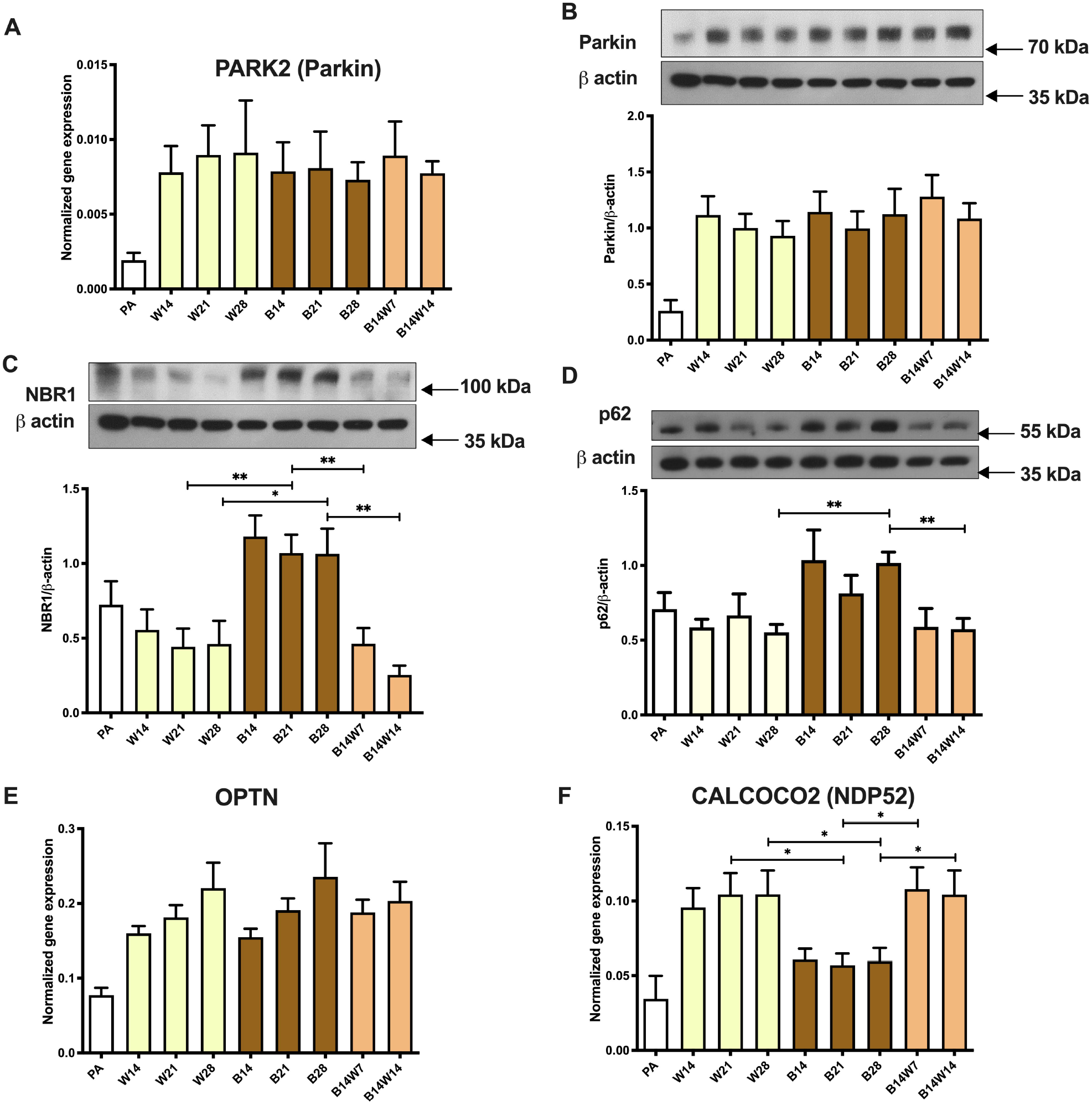
Expression of parkin-dependent mitophagy markers in white (W), beige (B) adipocytes, and in response to transition. Human primary abdominal subcutaneous preadipocytes were differentiated as in Figures 1-4. Quantification of parkin (A) gene and (B) protein expression. Protein expression of (C) NBR1 and (D) p62. Gene expression of (E) OPTN and (F) NDP52. Gene expressions were normalized to GAPDH and protein expressions to β-actin. Data are presented as Mean ± SD. n=6. *p<0.05, **p<0.01. Statistics: one-way ANOVA with Tukey’s post-test.

Next, we assessed the expression of other marker genes related to the adapter and parkin-dependent mitophagy pathway, OPTN (Figure 5E) and CALCOCO2/NDP52 (Figure 5F). Both genes were expressed at a low level in preadipocytes. OPTN was expressed at the same extent in white and beige adipocytes, and the transition did not influence its mRNA level (Figure 5E). CALCOCO2/NDP52 expression was increased during white adipogenesis and transition compared with the fully differentiated beige adipocytes (Figure 5F); this suggests the possibility of enhanced removal of the mitochondrial mass by the NDP52-dependent pathway.

### 2.5. Parkin-Independent Mitophagy-Related Genes Are Induced During Transition

Finally, to study whether parkin-independent mitophagy may contribute to beige to white transition, we investigated the expression of several genes that are involved in this pathway, FUNDC1, BNIP3, BNIP3L/NIX, FKBP Prolyl Isomerase 8 (FKBP8), and BCL2 Like 13 (BCL2L13) (Figure 6).The aforementioned markers were expressed at a low extent in undifferentiated progenitors. Similar expression levels of FUNDC1 and BNIP3 were found during white or beige differentiation. After four weeks of differentiation, the expression of BNIP3L/NIX, FKBP8, and BCL2L13 was repressed in beige compared to white adipocytes. Two weeks following the replacement of beige protocol to white, the investigated parkin-independent mitophagy markers were significantly upregulated (Figure 6). This data suggest that the parkin-independent pathway can also play an important role during beige to white transition of human subcutaneous adipocytes.

**Figure 6.**
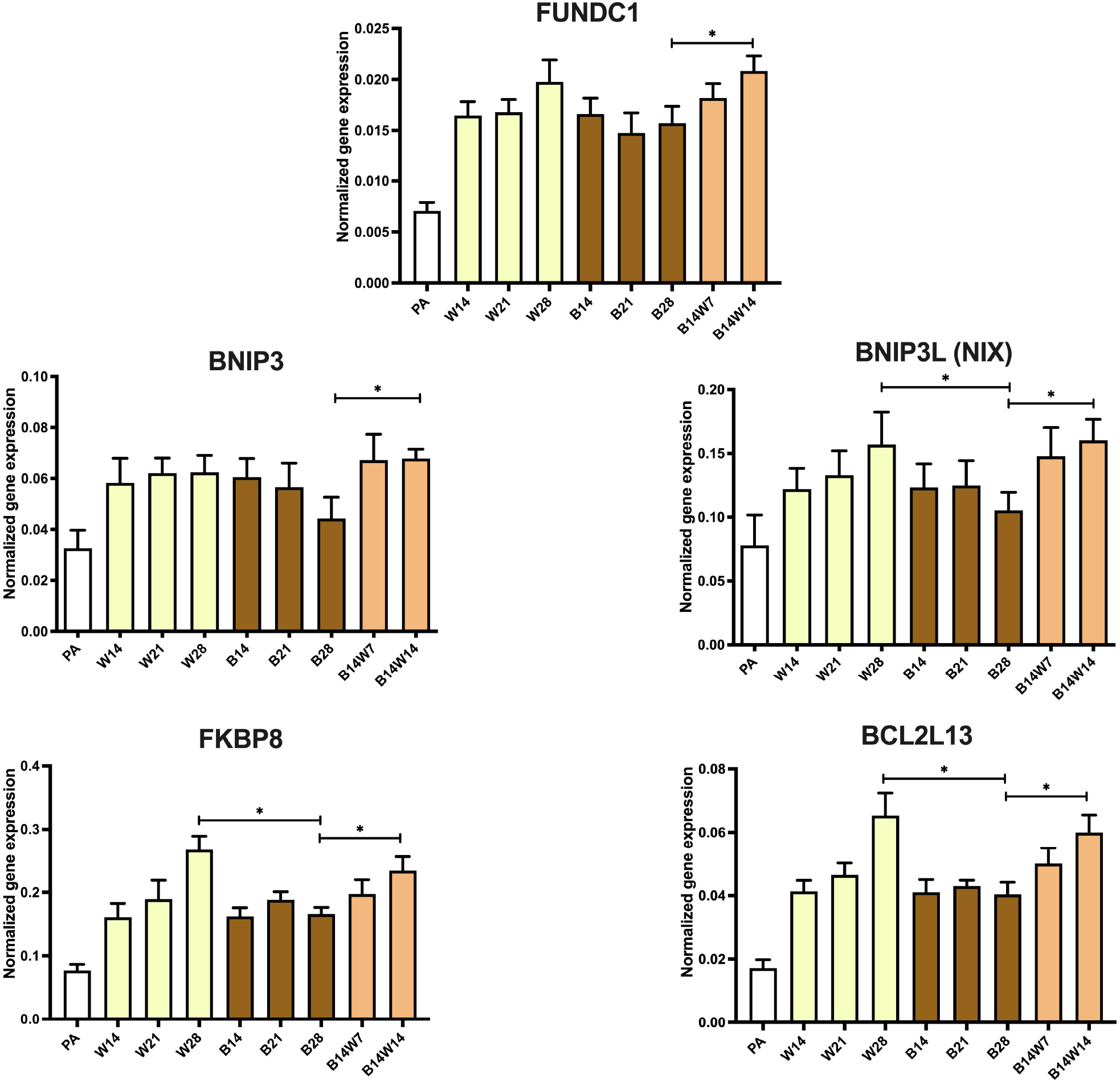
Expression of parkin-independent mitophagy markers in white (W), beige (B) adipocytes, and in response to transition. Human primary abdominal subcutaneous preadipocytes were differentiated as in Figures 1-5. Quantification of gene expression for FUNDC1, BNIP3, BNIP3L, FKBP8, and BCL2L13. Gene expressions were normalized to GAPDH. Data are presented as Mean ± SD. n=6. *p<0.05. Statistics: one-way ANOVA with Tukey’s post-test.

## 3. Discussion

BAT plays a central role in the energy homeostasis of mammals that are constantly exposed to cold challenge [12]. Following the detection of active BAT depots by nuclear imaging approaches in adult humans [55-57], a very strong negative correlation between obesity and the amount of active BAT was revealed [58,59]. Furthermore, the abundance of these depots strongly decreased in response to aging [60]. Independent studies have suggested that BAT depots in adult humans are predominantly composed of beige cells [13,61,62]. Transplants of human beige adipocytes improved diet-induced obesity and systemic metabolism in mice, which highlights the possibility of therapeutic application of beige cell implantation in the treatment of obesity and metabolic syndrome [63,64]. This inspired researchers to characterize beige adipogenesis and thermogenic activation in distinct human cellular models. To our knowledge, however, majority of these *ex vivo* studies have covered the differentiation period only maximum up to two weeks. Of note, the potential application of beige adipocyte activation or transplantation assumes that the applied cells maintain their energy expenditure for a significant period of time. Although abdominal subcutaneous WAT of human adults is not highly enriched in thermogenic adipocytes [17], it contains progenitors that can give rise to beige cells [65,66]. Due to its relatively easy accessibility, hASCs isolated from stromal-vascular fractions (SVFs) of abdominal subcutaneous fat biopsies or aspirations are frequently used for research and regenerative medicine [67].

In this study, we have attempted to follow up white and beige adipocyte differentiation of primary abdominal subcutaneous derived hASCs for four weeks. The extension of the PPARγ-driven beige differentiation resulted in further upregulation of UCP1, both at mRNA and protein level, while in white adipocytes it was expressed constantly at a moderate level (Figure 1A, B). This phenomenon was completely reproduced by our research group in Simpson-Golabi-Behmel syndrome (SGBS) adipocytes [44], which cell line is an accepted and widely used model of human white and beige adipogenesis [68,69]. Consistent with our previous findings [43,44], the size and locularity of lipid droplets were also different between the two cell populations during the entire differentiation (Figure 1C, D). In beige and brown adipocytes, mitochondria are critical for thermogenesis and energy metabolism. Mitochondria are fragmented in response to an adrenergic cue in rodent brown adipocytes, contributing to support uncoupled respiration and enhanced energy expenditure [70,71]. Pisani et al. have shown that UCP1-positive human adipocytes contain mitochondria mostly with fragmented morphology [48]. Recently, we found that cAMP-driven thermogenic stimulation resulted in increased mitochondrial fragmentation in human masked and mature beige adipocytes, which were differentiated from the same progenitor populations for two instead of four weeks [39]. When we sustained the beige differentiation, more mitochondria were fragmented in contrast to white adipocytes, in which these dynamic organelles remained elongated (Figure 1E).

The regulation of mitochondrial biogenesis and clearance is important for energy homeostasis and maintaining the optimal amount of mitochondria [34]. Mitochondrial biogenesis is controlled by several nuclear-coded transcriptional regulators, such as PGC1α [72]. Although the expression of PGC1α at mRNA tended to be elevated at the early phase of beige differentiation, it did not differ between white and beige adipocytes at a statistically significant level (Figure 2A). However, the amount of mtDNA showed an increasing tendency during the long-term beige adipogenesis (Figure 2B). The functional extracellular flux assay detected high basal, cAMP stimulated, and proton leak OCR and more prominent extracellular acidification, both at basal and activated conditions, in the case of beige adipocytes that were differentiated in the presence of rosiglitazone for four weeks. Respiration and extracellular acidification were significantly repressed in white adipocytes; however, they could be effectively stimulated in response to the cell permeable cAMP analogue, suggesting that some of the adipocytes that were differentiated in the presence of white cocktail are masked beige cells (Figure 2C, D).

Elevated mitochondrial fragmentation, mtDNA amount, and OCR raise the possibility of suppressed mitophagy in beige adipocytes. When the same progenitors were differentiated for two weeks, irrespective of the applied protocol, a few hours long cAMP treatment not only upregulated thermogenesis-related genes but also quickly downregulated mitophagy via protein kinase A, resulting in more mitochondria and increased UCP1 levels [39]. In the long-term differentiation settings, sustained rosiglitazone administration also resulted in a moderate suppression of mitophagy, shown by attenuated LC3-I to LC3-II conversion (Figure 3D), appearance of LC3 positive punctae (Figure 4B), co-localization of punctae and mitochondria (Figure 4C), and degradation of adapter proteins (Figure 5C, D). More research is needed to explore the underlying molecular mechanisms of how mitophagy is kept at a moderately low level in beige adipocytes.

Altshuler-Keylin et al. induced beigeing of WAT in male mice by intraperitoneal administration of the β3-adrenergic agonist, CL316243 for 7 consecutive days. Low autophagy activity was observed in the newly differentiated beige adipocytes [37], which is consistent with the ex vivo results presented here (Figures 3 and 4). After the withdrawal of this stimulus, they found that beige adipocytes lost their morphological and thermogenic characteristics and were converted to “white-like” adipocytes, which was triggered by mitochondrial clearance via mitophagy. During beige to white adipocyte transition, the expression of autophagy-related ATG5 and ATG12 genes were upregulated, the number of Green Fluorescent Protein (GFP)-LC3 puncta and the co-localization of GFP-LC3 and TOM20 were significantly increased, the protein level of LC3-II was elevated, and in parallel the selective autophagy adapter proteins, NBR1 and p62 were degraded, compared to the mice that were chronically treated with the β3-adrenergic agonist during the entire experimental period [37]. It has been shown in rodents that this adaptive transition process is mediated by selective autophagy without an intermediate precursor state, however, there are reports in which mature adipocytes from the mammary gland can de-differentiate at a certain level to progenitor-like cells then re-differentiate to adipocytes [73].

In our experiments presented here, we applied the human primary subcutaneous abdominal derived adipocyte *ex vivo* model to characterize the beige to white transition process in the context of mitophagy. UCP1 expression (Figure 1A, B), morphology of the lipid droplets (Figure 1C, D), mitochondrial fragmentation (Figure 1C, E), and basal respiration (Figure 2C) of the adipocytes were significantly altered as a result of the transition, therefore, these cells gained several characteristic features of the white adipocytes. A similar phenomenon was observed recently when we carried out 28 days differentiation in parallel with the replacement of beige protocol to white at the fourteenth day, on the SGBS preadipocyte line [44]. In contrast to our previous observations in SGBS adipocytes and the data presented here, Guennoun et al. observed a temporarily high UCP1-content of 14 days differentiated SGBS adipocytes even in response to the white differentiation protocol in the absence of any browning-inducers. Interestingly, when the white differentiation of SGBS cells was extended for two additional weeks, the expression of UCP1 was greatly declined [74]. The contribution of autophagy to this surprising finding has not been investigated so far.

Our data, shown in Figures 3 and 4, are consistent with the results of the *in vivo* study by the Kajimura group [37] and suggest that autophagy pathway is activated for the clearance of beige adipocyte mitochondria during the adaptive transition induced by the removal of browning-inducer, thereby regulating the entry of beige adipocytes into a thermogenically inactive dormant state. However, beige adipocytes that underwent transition *ex vivo* responded more effectively to the adrenergic stimulus mimicking dibutyril-cAMP by the activation of OCR, proton leak respiration (Figure 2C), and ECAR (Figure 2D) than the white ones. Of note, significant amount of UCP1 protein remained expressed after two weeks of transition (Figure 1B), which could underlie why converted adipocytes had higher stimulated and proton leak OCR compared to white ones. The observed functional differences suggest that converted beige and white adipocytes can be classified into two distinct cell populations. Systematic studies are needed to further explore the molecular signatures of the thermogenically active beige, converted beige, and white adipocytes in humans.

As the final step, we intended to clarify how the selective removal of mitochondria is mediated during the transition process. In the literature, contradictory data have been published about the involvement of parkin in the maintenance of murine beige adipocytes. During the adipogenesis of 3T3-L1 adipocytes, increased parkin expression was observed, whereas its expression was decreased as a result of rosiglitazone treatment [75]. Lu et al. found that parkin expression was induced during mouse beige adipocyte differentiation, moreover, they demonstrated the retention of mitochondria-rich beige adipocytes even after the elimination of adrenergic stimuli in PARK2 knockout mice [38]. In contrast to these findings, Corsa and colleagues found that parkin deletion in mouse adipocytes did not affect adipogenesis, beige to white transition, and maintenance of beige adipocytes [76]. Recently, we reported that parkin-dependent and independent mitophagy-associated genes were expressed in human masked and mature beige adipocytes, and the cAMP-driven thermogenic stimulus resulted in decreased expression of par-kin-dependent mitophagy-related genes [39]. In the current study, we have found that the gene and protein expression of parkin was not affected during beige to white transition (Figure 5A, B). However, the level of selective adapter proteins, NBR1 (Figure 5C) and p62 (Figure 5D), was significantly decreased and the expression of CALCOCO2/NDP52 (Figure 5F) and the investigated parkin-independent mitophagy-related genes (Figure 6) was significantly elevated during transition, compared to fully differentiated beige adipocytes. In summary, our data suggest that both parkin-dependent and independent mitophagy pathways are involved the regulation of mitochondrial elimination during beige to white adipocyte transition.

p62 is a multifunctional protein, involved in several signaling pathways affecting various cellular processes, such as inflammation, cell death, tumorigenesis, and metabolism [77,78]. It has been reported that whole-body p62 knockout mice show obese phenotype due to increased adiposity and reduced energy expenditure [79]. Furthermore, the mitochondrial function of BAT in adipocyte-specific p62^−/–^ mice was impaired, resulting in BAT to become unresponsive to β3-adrenergic stimuli [80].This suggests that p62 plays a significant role in the regulation of thermogenesis in BAT. A recent paper has demonstrated that NBR1 is required for the repression of adaptive thermogenesis via decreasing the activity of PPARγ in BAT of p62-deficient mice, thereby the inhibitory role of NBR1 in thermogenesis in the presence of p62 inactivation was identified [81]. Based on these studies, further investigations may reveal the role of p62 and NBR1 in the thermogenesis of human brown and beige adipocytes.

Individuals with obesity possess less active BAT but more “brownable” fat than the lean ones [17]. These “brownable” depots might contain large amount of beige adipocytes undergoing transition, in which autophagy and mitophagy are highly active. This is supported by the fact that ATG and autophagosome-related genes are highly expressed in visceral and subcutaneous WAT of patients with obesity [82,83]. In the future, well established molecular markers and histological or cell sorting methods are necessary to discriminate between white, active beige, and dormant beige adipocytes in distinct anatomical areas. This might allow researchers to analyze the gene expression changes during the conversion in individual cells that might reveal novel molecular targets, which control this process. A better understanding of the key molecular events that determine the entry into beige to white transition may offer new opportunities for specifically preventing this process in order to maintain active heat producing adipocytes; pharmacologically activated or transplanted for instance, in humans for improving energy metabolism and combatting obesity.

## 4. Materials and Methods

### 4.1. Materials

All chemicals were obtained from Sigma Aldrich (Munich, Germany) unless stated otherwise.

### 4.2. Isolation, maintenance, and differentiation of hASCs

hASCs were obtained and isolated from SVFs of human subcutaneous abdominal adipose tissue of healthy donors undergoing planned surgeries as previously described [39,43]. Absence of mycoplasma was ascertained by PCR Mycoplasma kit (PromoCell, Heidelberg, Germany). hASCs were seeded in 6 well plates (Costar, Corning, NY, USA) and differentiated for 14, 21, and 28 days as indicated, following the white [39,43,44] or beige [39,41,43,44] adipogenic differentiation protocols. White differentiation was initiated by using serum-free DMEM-F12 medium supplemented with 33 μM biotin, 17 μM pantothenic acid, 100 U/mL penicillin/streptomycin, 10 μg/mL human apo-transferrin, 20 nM human insulin, 200 pM triiodothyronine, 100 nM cortisol, 2 μM rosiglitazone (Cayman Chemicals, Ann Arbor, MI, USA), 25 nM dexamethasone, and 500 μM 3-isobutyl-1-methylxanthine (IBMX). Four days later, rosiglitazone, dexamethasone, and IBMX were removed. The beige protocol was initiated for four days, applying serum-free DMEM-F12 containing 33 μM biotin, 17 μM pantothenic acid, 100 U/mL penicillin/streptomycin, 10 μg/mL apo-transferrin, 0.85 μM human insulin, 200 pM triiodothyronine, 1 μM dexamethasone, and 500 μM IBMX. Then 500 nM rosiglitazone was added to the cocktail while dexamethasone and IBMX were omitted. Where indicated, after a 14 days beige differentiation period, transition was made to white differentiation protocol for further 7 or 14 days, respectively. Transition was initiated by addition of 100 nM cortisol and removal of 500 nM rosiglitazone, the key driver of beige differentiation. The concentration of human insulin was decreased by 42.5 times at the induction of transition. Media were replaced at an interval of 4 days.

### 4.3. Nucleic acid isolation, RT-PCR, and qPCR

Cells were collected in Trizol reagent (Thermo Fisher Scientific, Waltham, MA, USA) followed by manual isolation of RNA and DNA by chloroform extraction and isopropanol or ethanol precipitation, respectively. RNA quality was evaluated by Nanodrop (Thermo Fisher Scientific), and cDNA was generated by TaqMan reverse transcription reagent kit (Thermo Fisher Scientific) followed by qPCR analysis [39,84]. Gene expression was normalized to GAPDH. A list of all the probes is provided in Table S1. Quantification of mtDNA was performed by qPCR as previously described [39,46].

### 4.4. Antibodies and immunoblotting

Sample separation was performed by SDS-PAGE, followed by transfer to a PVDF membrane. The membrane was blocked by 5% skimmed milk solution [39,84]. The following primary antibodies were used: anti-UCP1 (1:750, R&D Systems, Minneapolis, MN, USA, MAB6158), anti-p62 (1:5000, Novus Biologicals, Centennial, CO, USA, NBP1-49956), anti-LC3 (1:2000, Novus Biologicals, NB100-2220), anti-Parkin (1:750, Santa Cruz Bio-technology, Dallas, TX, USA, sc-32282), anti-NBR1 (1:1000, Novus Biologicals, NBP1-71703), and anti-β-actin (1:5000, A2066). HRP-conjugated goat anti-rabbit (1:10,000, Advansta, San Jose, CA, USA, R-05072-500) or anti-mouse (1:5000, Advansta, R-05071-500) IgG were used as secondary antibodies. Immunoreactive proteins were visualized, followed by densitometry by FIJI as previously described [39].

### 4.5. Immunostaining and image analysis

hASCs were plated and differentiated in 8-well Ibidi μ-chambers (Ibidi GmbH, Gräfelfing, Germany) for the indicated number of days. Cells were washed once with PBS and fixed by 4% paraformaldehyde, followed by permeabilization with 0.1% saponin and blocking with 5% skimmed milk [39]. Primary antibody incubations were kept overnight with anti-TOM20 (1:75, WH0009804M1) and anti-LC3 (1:200, Novus Biologicals, NB100-2220). Secondary antibody incubation was kept for 3 hours with Alexa 647 goat anti-mouse IgG (1:1000, Thermo Fischer Scientific, A21236) and Alexa 488 goat anti-rabbit IgG (1:1000, Thermo Fischer Scientific, A11034). Propidium Iodide (1.5 μg/mL, 1 h) was used for nuclei labelling. Images were obtained by Olympus FluoView 1000 confocal microscope as previously described [39,46]. LC3 punctae and fragmented mitochondria were quantified using FIJI as previously described [39,46]. Co-localization of LC3 and TOM20 were evaluated by calculation of PCC [39]. Texture sum variance was calculated as described previously [43,45].

### 4.6. Determination of cellular OCR and ECAR

Cells were seeded and differentiated on XF96 assay plates (Seahorse Biosciences, North Billerica, MA, USA) and differentiated for 28 days with white, beige, or beige to white transition protocols, followed by measurement of OCR and ECAR with XF96 oximeter (Seahorse Biosciences). cAMP stimulated OCR, ECAR and stimulated proton leak OCR were measured as previously described [51]. Antimycin A (10 μM) was used for baseline correction. The OCR was normalized to protein content.

### 4.7. Statistics and figure preparation

All measured values are expressed as Mean ± SD for the number of independent repetitions indicated. One-way ANOVA with Tukey’s post hoc test were used for multiple comparisons of groups. Graphpad Prism 9 was used for figure preparation and statistics.

## 5. Conclusions

Although the concept of beige to white adipocyte transition is not novel, the underlying transcriptional and cell biological changes have not been investigated in human primary cell models so far. To our knowledge, we could successfully model white and beige differentiation of human abdominal subcutaneous adipocytes up to 28 days and beige to white transition *ex vivo* for the first time. Beige adipocytes had elevated mitochondrial biogenesis, UCP1 expression, fragmentation, and oxygen consumption compared to white. In adipocytes that underwent beige to white transition, these parameters were similar to those observed in white cells. During transition, both parkin-dependent and independent mitophagy marker genes were induced. The direct functions of individual elements of the mitophagy machinery need to be investigated by experiments involving pharmacological inhibitors and gene silencing, deletion, or overexpression. If pharmacologically stimulated and/or transplantable beige adipocytes are available for combatting obesity, inhibition of beige to white transition might be an additional approach to maintain their high energy expenditure. This potentially can lead to more amount of active BAT instead of “brownable” fat.

## Supplementary Materials

Figure S1: Secondary antibody control images showing the specificity of the antibodies used for LC3 and TOM20 immunostaining, Table S1: Table listing all gene expression assays used in the study.

## Author Contributions

Conceptualization, E.K. and M.S.-T.; methodology, Z.Bac.; software, A.S., G.M., and Z.Bac.; validation, E.K. and M.S.-T.; formal analysis, A.V., A.S., G.M., and Z.Bac.; investigation, A.V., A.S., K.V., I.C., and M.S.-T.; resources, Z.Bal. and C.L.; data curation, A.S. and M.S.-T.; writing—original draft preparation, A.V., A.S., and M.S.-T.; writing—review and editing, E.K.; visualization, A.S.; supervision, E.K.; project administration, E.K. and M.S.-T.; funding acquisition, E.K. All authors have read and agreed to the published version of the manuscript.

## Funding

This research was funded by the National Research, Development and Innovation Office of Hungary, NKFIH-FK131424 and the European Union and the European Social Fund, EFOP-3.6.3-VEKOP-16-2017-00009. The APC was funded by NKFIH-FK131424.

## Institutional Review Board Statement

The study was conducted according to guidelines of the Declaration of Helsinki and approved by the Medical Research Council of Hungary (20571-2/2017/EKU; 28.04.2017). Experiments were performed strictly in accordance with the approved ethical regulations and guidelines.

## Informed Consent Statement

Written informed consent was obtained from all participants prior the collection of tissue samples.

## Data Availability Statement

The data presented in this study are available on request from the corresponding author.

## Acknowledgments

We thank Dr. László Fésüs for his continuous mentoring, consultation, and for reviewing the manuscript. We acknowledge Jennifer Nagy for excellent technical assistance.

## Conflicts of Interest

The authors declare no conflict of interest. The funders had no role in the design of the study; in the collection, analyses, or interpretation of data; in the writing of the manuscript, or in the decision to publish the results.

## Notes

### Competing Interest Statement

The authors have declared no competing interest.

