## Supplementary Figures and Tables for "Mitophagy mediates beige to white transition of human primary subcutaneous adipocytes *ex vivo*"

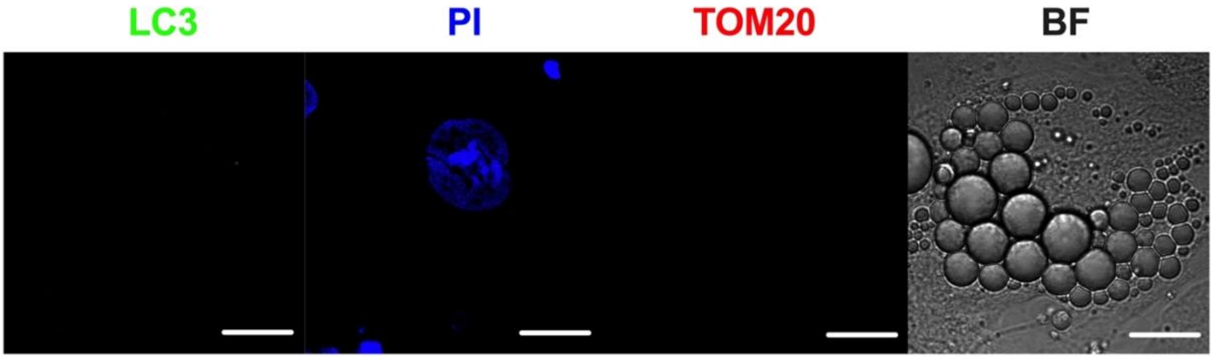


**Figure S1: Secondary antibody control images showing the specificity of the antibodies used for LC3 and TOM20 immunostaining.** PI labels the nucleus. BF represents brightfield image. Scalebars represent 10μm.

**Table S1: Table listing all gene expression assays used in the study**

| Gene name | Assay ID |
|---|---|
| UCP1 | Hs00222453_m1 |
| PPARGC1A | Hs01016719_m1 |
| GAPDH | Hs99999905_m1 |
| PARK2 | Hs01038322_m1 |
| SQSTM1 | Hs00177654_m1 |
| OPTN | Hs00184221_m1 |
| NDP52 | Hs00977443_m1 |
| BNIP3 | Hs00969291_m1 |
| BNIP3L | Hs00188949_m1 |
| FKBP8 | Hs01014664_m1 |
| BCL2L13 | Hs00209789_m1 |
| FUNDC1 | Hs00697693_m1 |
